# Deep Learning-based Microbubble Localization for Ultrasound Localization Microscopy

**DOI:** 10.1101/2022.02.02.478911

**Authors:** Xi Chen, Matthew R. Lowerison, Zhijie Dong, Aiguo Han, Pengfei Song

## Abstract

Ultrasound localization microscopy (ULM) is an emerging vascular imaging technique that overcomes the resolution-penetration compromise of ultrasound imaging. Accurate and robust microbubble (MB) localization is essential for successful ULM. In this study, we present a deep learning (DL)- based localization technique that uses both Field-II simulation and *in vivo* chicken embryo chorioallantoic membrane (CAM) data for training. Both radiofrequency (RF) and in-phase quadrature (IQ) data were tested in this study. The simulation experiment shows that the proposed DL-based localization was able to reduce both missing MB localization rate and MB localization error. In general, RF data showed better performance than IQ. For the *in vivo* CAM study with high MB concentration, DL-based localization was able to reduce the vessel MB saturation time by more than 50% as compared to conventional localization. Additionally, we propose a DL-based framework for real-time visualization of the high-resolution microvasculature. The findings of the paper support the use of DL for more robust and faster MB localization, especially under high MB concentrations. The results indicate that further improvement could be achieved by incorporating temporal information of the MB data.

## I. Introduction

AS a diffraction-limited imaging modality, the performance of ultrasound imaging has long been limited by the classical trade-off between imaging resolution and penetration depth. For decades, there has been a long quest to break the diffraction limit of ultrasound [1]. For example, various attempts had been made to super-focus the acoustic beams through negative refraction generated by acoustic metamaterials [2] [3] [4]. In a medium with random and inhomogeneous echogenicity, the diffraction limit may be overcome by retransmission and refocusing of received signal through a time reversal mirror [5]. As an acoustic analogy to structured illumination microscopy (SIM), acoustical structured illumination was recently proposed to surpass the resolution limit by generating a series of known patterns with the transducer, which enables encoding of high-resolution information of the observed image [6]. The lateral resolution of ultrasound can also be improved by null subtraction imaging (NSI), a technique that applies multiple receive apodizations to reduce sidelobes and enhance the mainlobe [7]. Superresolution of these techniques is achieved contrast-free and based on manipulation of the transmit and/or received point spread function (PSF) of the ultrasound imaging system.

Recently, a contrast microbubble (MB)-based technique named ultrasound localization microscopy (ULM) [8] [9] was proposed to overcome the diffraction limit of ultrasound based on the principle of localization microscopy (e.g., PALM [10] [11] and STORM [12]). MBs have a similar size to red blood cells, and they travel within the vasculature for several minutes post intravenous injection [13]. A super-resolved blood vessel density map can be constructed by localizing and accumulating the MB locations in a consecutive sequence of diffraction-limited image frames. Moreover, a high-fidelity blood flow speed map can be recovered from measuring the frame-to-frame displacement of MBs. Various studies have demonstrated multiple applications of ULM under clinical and preclinical settings, including tumor characterization [14] [15] [16], brain imaging [17], and abdominal organs such as liver, kidney, and pancreas [18].

At present, the clinical potential and utility of ULM are challenged by various limitations of ULM, including the time-consuming imaging acquisition, the high computational cost of post-processing steps, the lack of ground truth to validate ULM *in vivo*, and other practical challenges such as tissue motion caused by breathing and scanning. These limitations and challenges remain active topics of research. In this study, we focus on addressing the issues of MB localization that contribute to the time-consuming data acquisition and high computational cost associated with post-processing. The current standard practice for MB localization is based on using an estimated template of the ultrasound system impulse response (i.e., the PSF) to identify individual MB signals and measure the physical location of the MB (e.g., by estimating the centroid of the MB signal). However, an ultrasound imaging system has a spatially-variant PSF that is subject to many confounding factors such as noise, phase aberration, multipath reverberation, and attenuation. Estimation of PSF is a challenging task, which leads to unreliable MB localization. In addition, as shown in [19] [20], MB localization based on the centroid of the MB signal is subject to errors induced by the non-linear MB response and the spatial sampling of the MB signal. Meanwhile, even with an ideal localization algorithm and MB data, the accuracy of MB localization is still subject to the Cramer-Rao lower bound, which is approximately one-tenth of the ultrasound wavelength [21]. Since MB localization occurs early in the ULM processing workflow, errors introduced in the localization step propagate downstream to affect the performance of subsequent operations, resulting in suboptimal ULM image quality. Therefore, improving the performance of MB localization is an essential task for improving the overall robustness of ULM.

Another challenge associated with MB localization in practice is the trade-off among MB concentration, data acquisition time, and localization accuracy. A diluted concentration of MB creates more MB signal separation in space, which makes the MB locations less ambiguous and easier to estimate. However, dilution of MB also slows down MB perfusion, resulting in longer acquisition times for full vascular reconstruction [22]. On the other hand, although high MB concentration leads to faster vessel MB saturation, the amount of spatially overlapping MB signals also substantially increases. This leads to a significant amount of wasted MB signals because overlapping MB signals are difficult to localize with conventional localization techniques and subsequently discarded. Although MB signals with partial overlaps can be potentially separated and localized, the overlap may distort the individual MB signal and challenge the pre-defined PSF. In reality, especially under clinical settings for human imaging, it is difficult to modulate the MB concentration in the bloodstream, which is largely dictated by physiology. In addition, the clinical dose of MB administration and MB concentration are typically regulated, which is difficult to change. Therefore, developing a robust MB localization method that works for various MB concentrations with various amounts of MB overlaps (especially for high MB overlap) is critical for the successful preclinical and clinical translations of ULM.

Several methods have been proposed to address the issue of high MB overlap at high MB concentrations. Zhang *et al*. proposed the use of phase change nanodroplets for flow and concentration independent localization microscopy with reduced acquisition time [23] [24]. Recently, we proposed a MB separation technique based on Fourier-based filtering to address the issue of MB overlap by decomposing high MB-count data into sub-sets of low MB-count data [25]. The MB separation was achieved by leveraging the different movement speeds and directions of MBs, which manifest as different slow-time frequency components in the Fourier space. The method greatly improved the amount of MB signals that can be localized and subsequently the overall quality of ULM imaging. However, the assumption of utilizing temporal frequency difference to separate MB signals may break down when MB concentration is high and/or flow hemodynamics are complex. Also, by separating the data into subsets with identical dimensions as the original data, the MB separation technique exacerbates the issue of the high computational cost of ULM. Therefore, it may be challenging in practice to implement the MB separation technique, and a faster and better MB localization method remains to be developed.

Considering ongoing breakthroughs in deep learning (DL) and computer vision, DL-based techniques have gained popularity in medical image processing, especially for tasks requiring a high level of abstraction [26]. In ultrasound imaging, DL implementation has mostly been focused on improving beamforming [27] [28] [29] [30] [31]. For DL-based ULM, van Sloun *et al*. [32] and Liu *et al*. [33] validated the feasibility of DL-based MB localization using different classes of neural network architectures. Both studies have shown promising potential for applying DL for accelerated and robust MB localization. Liu *et al*. also used *in vivo* vessel networks to generate more realistic training data. Youn et al. [34] performed convolutional neural network (CNN) -based localization on RF channel data generated by Field-II simulation, which showed improvement over local peak detection using envelope-detected signals on both simulation and phantom data. Lok *et al*. [35] used data from chicken embryo chorioallantoic membrane (CAM) for *in vivo* validation with optical images as the reference. They showed that DL-based MB localization outperformed conventional localization techniques under low to moderate MB concentrations. As an alternative to the common localize-and-track workflow for ULM, Milecki et al. [36] proposed a DL-based framework that recovers tracks directly from MB signal without explicitly localizing individual MBs. Since velocity information is not retained in the recovered tracks, this alternative framework is suited for applications where blood flow dynamics information is required. In our study, we will focus on the typical ULM framework with explicit localization of MBs.

Different from existing studies where low to moderate MB concentrations were typically used to develop DL-based MB localization, our objective in this study was to develop and test the performance of DL-based MB localization under high to very high MB concentrations. The advantage of using high MB concentration is the faster MB perfusion speed in tissue microvasculature, which shortens ULM data acquisition time that is significant for clinical implementations of ULM. To the best of our knowledge, whether DL-based localization can withstand the high MB concentration scenario is yet to be explored. In this study we characterized the performance of various MB localization techniques under a wide range of MB concentrations by using Field-II ultrasound simulations as well as vascular graph models extracted from *in vivo* data for training the neural network. Our training data consists of realistic vascular structures extracted from optical images of chicken embryo chorioallantoic membrane (CAM) surface vessels. We also explored the use of RF data to investigate localization with both amplitude and phase information. Our method was applied to several different scenarios: simulated test data ranging from very sparse to very dense MB distributions, for the characterization of how the performance gain of DL-based MB localization scales with increasing MB concentration; simulated test data of high concentration MB within vascular structure, for the study of the potential reduction in acquisition time using DL-based localization; and on *in vivo* CAM data with high concentration MB injection and high-resolution optical images as the ground truth. In addition to using DL-based MB localization for conventional ULM processing, we also proposed a high-resolution blood flow visualization method using DL-processed B-mode ultrasound MB data to demonstrate the feasibility of potential real-time ULM imaging.

## II. Methods

### A. Field-II simulation for MB data synthetization

To study the effect of MB distribution in the training data, three groups of training data sets were generated based on Field-II [37] [38] and used to train the networks: group 1 uses low MB concentration of up to 300 MBs/mm^2^ (1.77MBs/λ^2^) Group 2 and 3 both have higher MB concentration closer to *in vivo* experiments, where group 2 uses randomly distributed MB signal (in space) for training and group 3 uses the CAM vessel model for assigning MB positions. The hypothesis was that the added information of microvascular structure (e.g., vessel vs. non-vessel regions, large vs. small vessels) would facilitate DL performance in MB localization.

For each training group, we generated 9,000 training images and 1,000 validation images of MBs within a 1 mm × 1 mm field-of-view (FOV), which corresponds to 203 × 203 pixel images with a 4.928-μm pixel size. This FOV was selected to balance several factors including being able to capture structural features of the vessel network on multiple scales, to not exceed the computer memory constraints, and to maintain the efficiency of training the neural network. With confocal imaging [39], we were able to observe ~25 MBs in a 200 μm × 200 μm FOV, which translates to ~625 MBs/mm^2^. Therefore, we used a normal distribution with μ = 600, σ = 600 MBs/mm^2^ to regulate MB concentrations in our high concentration simulations (group 2 and 3).

For the group 1 and group 2 training set, the axial and lateral coordinates of the MBs were drawn from a uniform distribution within the dimension of the imaging FOV. For the group 3 training set, optical images (see Section II-D for optical imaging details) of the CAM surface vessels were used to place MBs in the vascular architecture (Fig. 1). Binary vessel maps were generated from the green channel (which provides the best contrast for vessels) of each optical image using MATLAB’s adaptthresh function, which computes local threshold based on Gaussian weighted mean of neighboring pixels (Fig. 1b). A subregion (1 mm x 1 mm) within the binary maps was randomly selected for the simulation of each training image. Training images with different MB concentrations were generated in a similar way as in group 1 and 2.

**Fig. 1.**
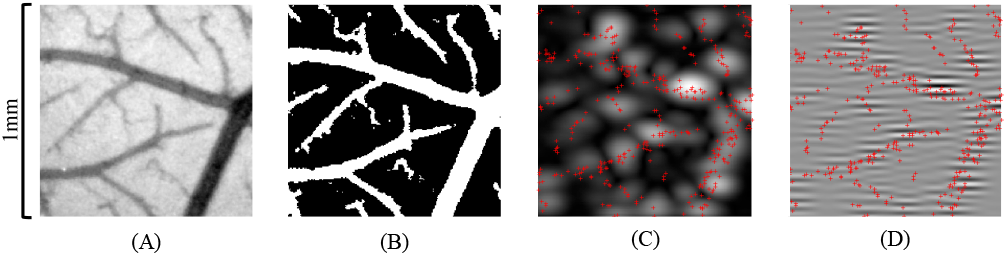
Example of Group 3 training data generation based on optical imaging of the CAM. (A) Green channel image of the CAM surface. (B) Binary vessel map after adaptive thresholding. (C) and (D) are simulated B-mode (i.e., envelope detected) image and RF data using Field II, respectively. Red crosses mark the true MB locations.

The simulation sequence was configured according to the CAM ultrasound imaging settings described in Section II-D. Details of the simulation specifications are summarized in Table I. Simulated ultrasound data were interpolated to the resolution of 4.928 μm. Finally, Gaussian white noise with a level of 10±2% of the ultrasound signal peak amplitude was added to the ultrasound data. The noise level was selected based on measurements from our experimental CAM data.

**Table I.** Field-II Simulation Parameters

| Variable name | Value |
| --- | --- |
| Transducer type | Linear array transducer |
| Number of transmit elements | 128 |
| Number of receive elements | 128 |
| Element height | 80 μm |
| Element width | 62 μm |
| Pitch | 70 μm |
| Transmit apodization | Kaiser |
| Receive apodization | Rectangle |
| Center frequency | 20 MHz |
| Sampling frequency | 125 MHz |
| Speed of sound | 1540 m/s |
| Number of compounding angles | 9 |

### B. Deep learning (DL) model design

U-Net is a CNN architecture first proposed in 2015 for biomedical image segmentation [40]. The architecture and its variations have since been successfully applied to many different tasks in biomedical image processing. Here, we adopt the encoder-decoder structure of the U-Net for our CNN model. The model contains a feature extraction path (the encoder) that extracts high-dimensional feature maps from the input ultrasound data, and a reconstruction path (the decoder) that extracts MB locations from the embedded feature map.

Fig. 2 is a schematic diagram of the network. Each downsampling block contains two convolution-batch normalization-activation units. Each reconstruction block starts with a 2 × 2-kernel convolution that halves the number of channels. The output of the corresponding feature extraction block is upsampled to match the spatial dimension of the output of the 2×2 convolution layer before they are stacked along the channel axis. The stacked data then go through convolution-batch normalization-activation units and are up-sampled by a factor of 2. Dropout is implemented in the bottleneck layer to prevent overfitting [41]. The final output block contains two 2-D convolution layers. The network contains a total of 8 convolution layers in the encoder part, and 8 convolution layers in the decoder part. We used leaky ReLU as the activation function, defined as:

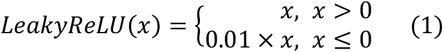

where *x* is the output of a convolution layer after batch-normalization. The model was trained with the Adam optimizer with a learning rate of 0.001 [42]. The loss function was defined as:

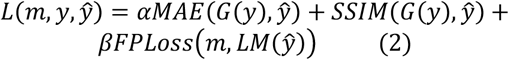

where *y* is the ground truth binary localization map, *m* is the ground truth binary vessel mask, 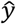 is the predicted localization map, *G* is a Gaussian filter with *σ* = 2 pixels (pixel size is 4.928 μm), and *MAE* is the mean absolute error, defined as

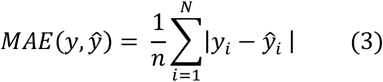

where *y_i_* and 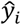 denote the i^th^ entry in the ground truth and the prediction, respectively. SSIM represents the structural similarity [43]:

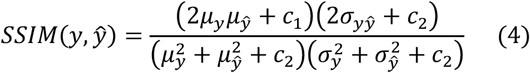

where *μ_y_*, *σ_y_* are the average and the standard deviation of the pixel values in ground truth image, 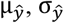 are the average and the standard deviation of the pixel values in the prediction, and 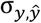 is the covariance of the ground truth and the prediction. *c*_1_ and *c*_2_ are the stabilizer terms in case of division by a very small denominator, defined as

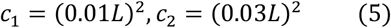

where *L* is the dynamic range of the image. *FPLoss* is a term that penalizes false positive predictions (i.e., false MB localizations), defined as the number of MB localizations that are outside of vessels, divided by the total number of predicted positives:

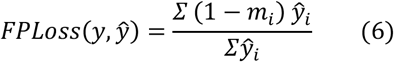

where *m*_i_ is the *i^th^* entry in the binary ground truth vessel map. Since *FPLoss* is a metric for binary prediction, and the output of the network is not necessarily binarized, a local maximum filter *LM* was used to convert the output to a binary localization map where the pixels of the local peaks were set to 1. *α* and *β* are weights for the *MAE* and *FPLoss* terms, respectively. *α* was set to 0.001 for all training sets. *β* was set to 1 for training sets that contain vessel structure and 0 for training sets without vessel structure. The loss function was designed so that the main objective function to reduce was still the SSIM. The addition of the *MAE* term helps the model to better preserve the intensity of the produced output, while *FPLoss* provides the network with additional information regarding the vascular structure, that is not explicitly given by the map of the true MB locations. False MB localization within nonvascular regions will be penalized heavier than false MB localization within vascular regions.

**Fig. 2.**
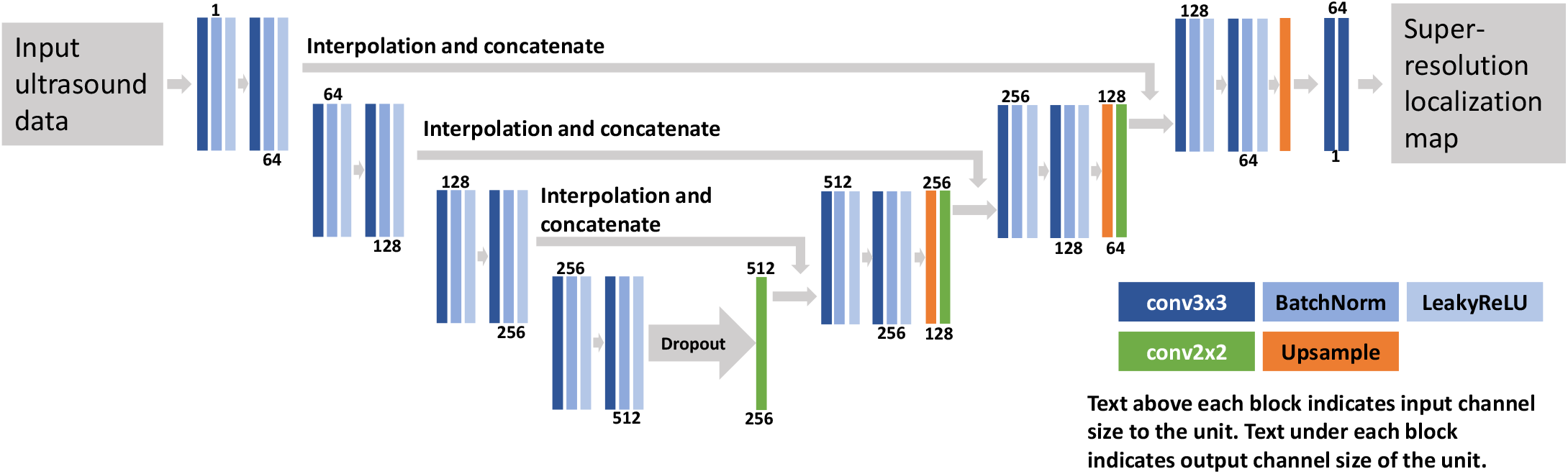
Schematic diagram of the neural network. The color-coded blocks represent different types of layers in the NN. The NN contains a feature-extraction path that extracts feature maps from the input ultrasound data, and a reconstruction path that recovers a super-resolution localization map from the feature maps

After each epoch of model update using the 9,000 training images, the training performance was evaluated on the validation set of 1,000 images. Hyper-parameter selection relies mostly on the validation set performance. Training samples were fed into the network as mini-batches of 16 images. The final trained model generates “sharpened” MB signals (Fig. 3), reducing the dimension of the apparent PSF size, where overlapping MBs signals in the original input are now separated.

**Fig. 3.**
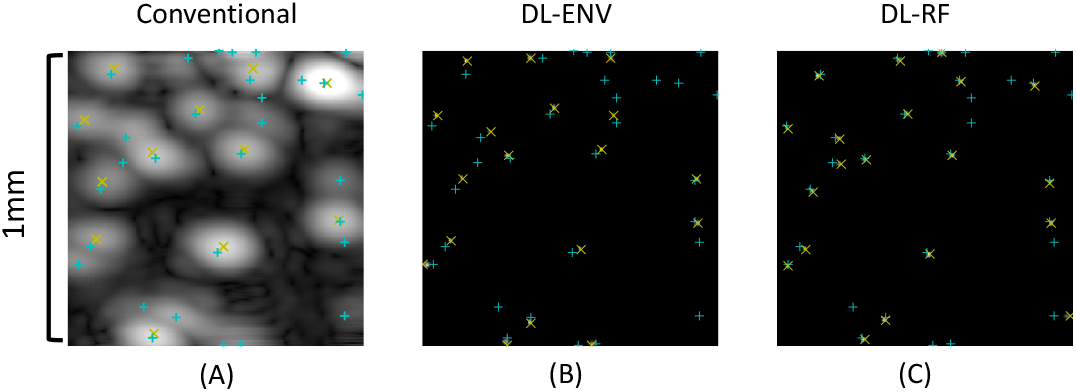
(A). Simulated B-mode (envelope detected) image of the MBs and the results of conventional localization. (B). Output of the model trained with envelope detected (ENV) data. (C). Output of the model trained with RF data. In all subfigures, measured locations are marked by yellow × and true MB locations are marked by cyan +. DL enhances the input ultrasound data by sharpening the MB signal so that they can be better separated for easier localization.

### C. ULM data processing

For all ULM data except simulation, the tissue signal was first removed by singular value decomposition (SVD)-based clutter filtering [44]. For MB localization with conventional, non-DL MB images, a cross-correlation-based approach as described in [45] was used with a synthesized multivariate Gaussian PSF. For the DL-processed MB data, the location of each MB was extracted by the blob detection function available in the scikit-image Python package using the difference-of-gaussian method [46]. The extracted locations were accumulated across each imaging frame to generate the microvessel density map. To better evaluate the performance of MB localization across different methods, no MB pairing or tracking (i.e., raw accumulation only) was implemented in our processing.

In addition to conventional ULM processing, the DL-enhanced MB data can also be used for real-time, high-resolution blood flow display: a sliding window of size *W* in temporal direction was used to directly accumulate the DL-enhanced MB data to display blood vessels with high spatial resolution. A matrix keeps the latest *W* DL-processed frames of MB data. When a new frame is processed by the network, the matrix will be updated by removing the oldest frame with the new frame. A new frame of output display was obtained by summing all the *W* frames of the updated matrix in the time direction.

### D. In vivo ULM data acquisition

Fresh fertilized chicken eggs were provided by the University of Illinois Poultry Research Farm and housed in a tilting incubator (Digital Sportsman Cabinet Incubator 1502, GQF Manufacturing Inc.). After four days, the eggshells were removed and embryos were transferred into plastic weigh boats. The *ex ovo* CAMs were then placed into a humidified incubator (Darwin Chambers HH09-DA) until the day of imaging, 13 days after eggshell removal.

In preparation for contrast MB injection, a glass capillary needle was produced by pulling a borosilicate glass tube (B120-69-10, Sutter Instruments, Novato, CA, USA) with a PC-100 glass puller (Narishige, Setagaya City, Japan). The glass needle was attached to a 1-mL syringe using Tygon R-3603 laboratory tubing. The surface vasculature of the CAM was punctured with the glass capillary needle and 50 μL of the Definity solution (Lantheus Medical Imaging Inc.) was injected into the embryo immediately prior to imaging.

Optical images were acquired using a Nikon SMZ800 stereomicroscope (Nikon, Tokyo, Japan) with a mounted DS-Fi3 digital microscope camera (5.9-megapixel CMOS image sensor, Nikon). Optical data were recorded using the Nikon NIS-Elements software platform and exported for offline analysis.

Ultrasound data were acquired using a Vantage 256 system with an L35-16vX high-frequency linear array transducer (Verasonics Inc., Kirkland, WA). The transducer was placed on the side of the plastic weigh boat to image the surface vasculature of the CAM. Imaging was performed with a center frequency of 20 MHz, using 9-angle plane wave compounding (1-degree increments) with a post-compounding frame rate of 1,000 Hz. Ultrasound data was saved as in-phase quadrature (IQ) datasets of 1,600 frames each for a total acquisition length of 32 seconds (32,000 frames). RF data was obtained from IQ data based on IQ demodulation [47].

## III. Results

### A. Individual MB localization performance for low to moderate MB concentration

The performance of the DL-based localization was quantified on a set of Field-II simulation data with low to moderate MB concentration using models trained with group 1 low concentration data. The evaluation focus on the ability to correctly identify individual MB signal. The testing set consists of 1mm×1mm samples with MB concentration ranging from 1 MB/mm^2^ to 100 MBs/mm^2^ (0.0059-0.59 MBs/ λ^2^), 50 samples were generated for each concentration. DL-based localization was performed on each of the simulated data. The DL localized MB locations and true MB locations were paired by using the algorithm proposed in [45]. True MB locations that were not paired, as well as paired true MB locations where the pairwise distance was greater than a threshold *tol*, were considered false negatives (FN_MB_). Similarly, predicted MB locations that were not paired, as well as paired predicted MB locations where the pairwise distance is greater than *tol*, were considered false positives (FP_MB_). The remaining paired MB locations were considered true positives (TP_MB_) and subject to localization error measurement.

An evaluation metric that includes three criteria were developed to quantitatively measure MB localization performance: MB false discovery rate (FDR), MB miss rate, and mean localization error, defined as:

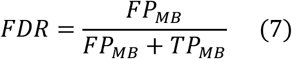

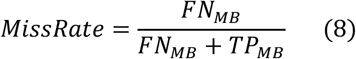

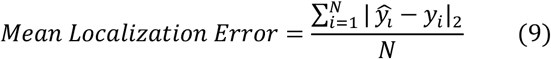

where 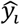 is the measured true positive MB locations, and *y_i_* is the corresponding ground truth. The performance of ENV-trained network (DL-ENV), RF-trained network (DL-RF), and conventional localization were averaged across all the 50 testing images for each MB concentration. Fig. 4 plots the average performance of DL based localization and conventional localization against increasing MB concentration using the three metrics described above with *tol* set to 5 pixels (0.32λ). The shaded regions represent the standard deviation of the performance within each concentration. Both DL-based localization methods outperformed conventional localization under this setting. DL-RF outperforms DL-ENV in all criteria except for the low concentration FDR, which can be a result of DL-RF having the overall tendency of localizing more MBs than DL-ENV. Conventional localization has particularly poor performance compared to DL-ENV and DL-RF for low concentration in FDR because it lacks the ability to distinguish noise and artifact from actual MB signal. The gap between DL-ENV and DL-RF increases in the mean localization error and the false discovery rate as concentration increases, which indicates that the addition of phase information in the RF data helps to better maintain robustness of localization as the amount of overlapping MB signal increases.

**Fig. 4.**
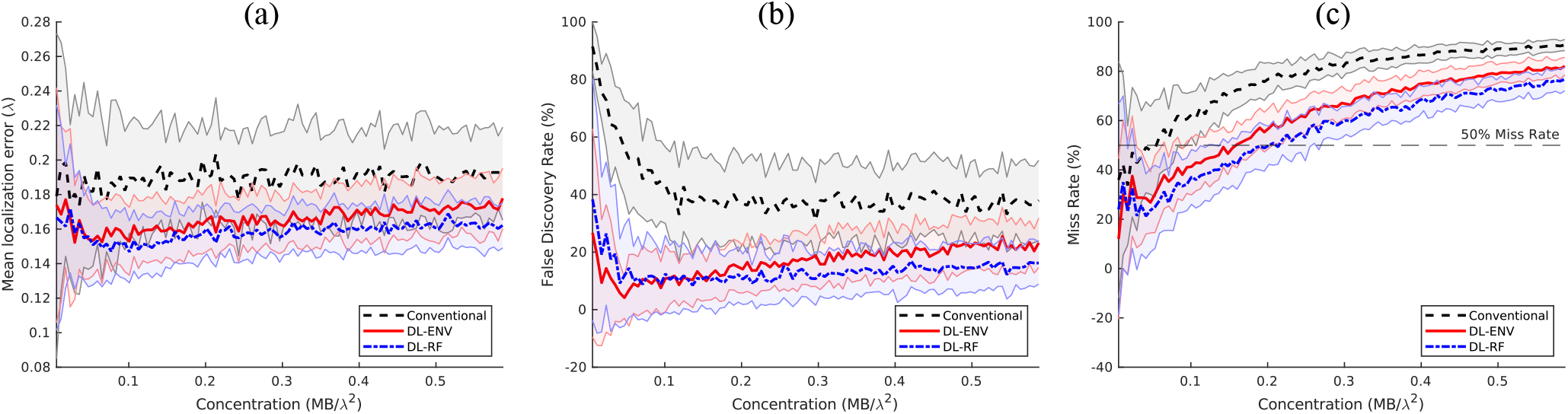
Average localization performance on the low concentration simulation test dataset.

We are specifically interested in the case of missing localizations, as it is closely related to whether increasing MB concentration is still profitable in terms of reducing the acquisition time required for full reconstruction of the vascular structure. We identify the point where a localization method hits above 50% miss rate as the indicator that it stops effectively localizing individual MBs. The 50% miss rate was marked on the performance graph as horizontal dashed lines. Fig. 5 plots the concentrations at which each localization method hits above 50% miss rate against different *tol* value chosen to define a missed localization (FN_MB_). Both figures showed that DL-based localization can withstand higher concentration before accurate localization of individual MB becomes a task too challenging, which can be useful for data acquired with high MB concentration, as well as for complex or hierarchical vasculature, where large vessels are present in the imaging plane. Specifically, DL-RF localization is able to maintain less than 50% miss rate using 4 times higher MB concentration compared to conventional localization.

**Fig. 5.**
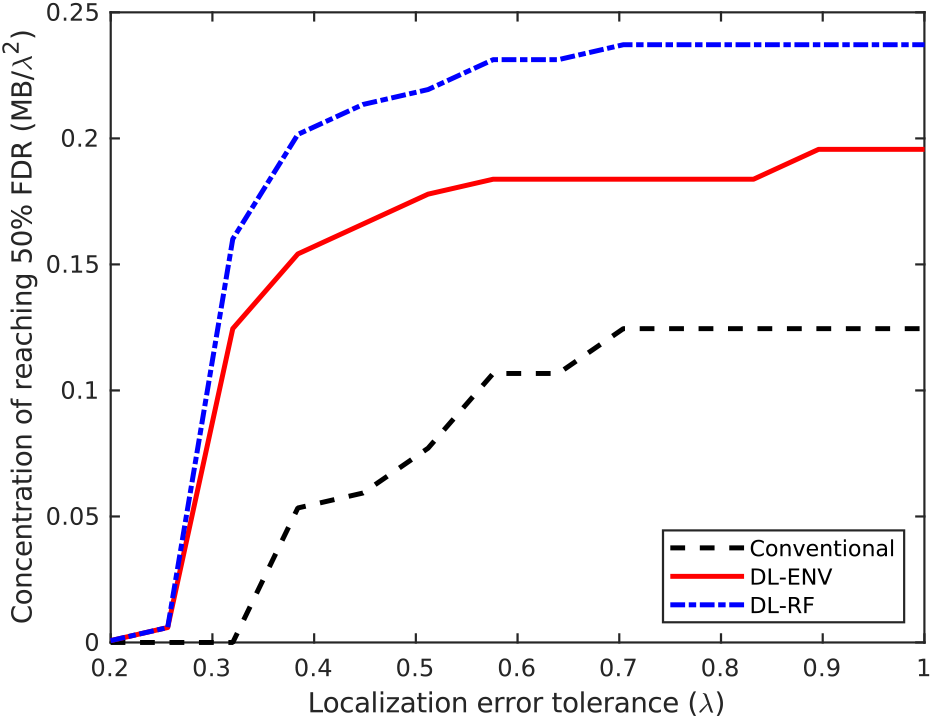
The point where each localization method hits above 50% miss rate versus different tolerance value for localization error for the simulation dataset.

However, reaching the “breakpoint” for individual MB localization does not necessarily mean the complete breakdown of ULM algorithm. Localization of the centroids of “blobs” of MBs can still provide useful information for reconstruction of the vascular structure. Moreover, as the blobs often tend to stay clustered for several consecutive frames, motion tracking on the entire blobs can still be used for estimation of flow speed. Therefore, we will further extend the concentration range to study the performance of each localization method under more extreme scenarios that are likely to be closer to real experimental environment.

### B. Individual MB localization performance for high concentration MB data

The concentration of simulation data was extended to 1000 MBs/mm^2^ (5.9MBs/ λ^2^) with a 10-MBs/ mm^2^ step size, with 50 samples generated for each concentration. The DL model was retrained using group 2 and group 3 high concentration training data. Evaluation of each method was carried out using the same method described in Section II.A. The behavior of the trained model changes with significantly increased MB concentration. Fig. 6 compares the performance of models trained with simulation with and without the CAM vessel structure against conventional localization. Fig. 7 compares the MB localization performance between conventional and DL-based methods based on CAM vessel training data only. Similar performance as the low to moderate concentration was observed for the extended concentration ranges. DL-based localization methods in general showed better performance than conventional localization across the board, with DL-RF showing the best localization performance overall. DL-based localization with the CAM vessel structure incorporated in simulation had better MB localization performance than without. FDR was significantly reduced, while minor improvements were observed for mean localization error and miss rate. The significantly lower false positives for both ENV and RF trained networks indicates that the spatial information associated with vessels was indeed helpful for the network to differentiate vessel versus non-vessel regions to facilitate better MB localization. DL-RF localization was able to reduce the MB miss rate while keeping the FDR at a similar level with conventional localization. This is not surprising because as hypothesized, RF data has both the amplitude and phase information of MBs, which facilitates MB separation and localization. The performance gain of DL-ENV localization was worsened compared to the low concentration case, which indicates that without phase information, the localization method is more sensitive to increased overlapping MB signal.

**Fig. 6.**
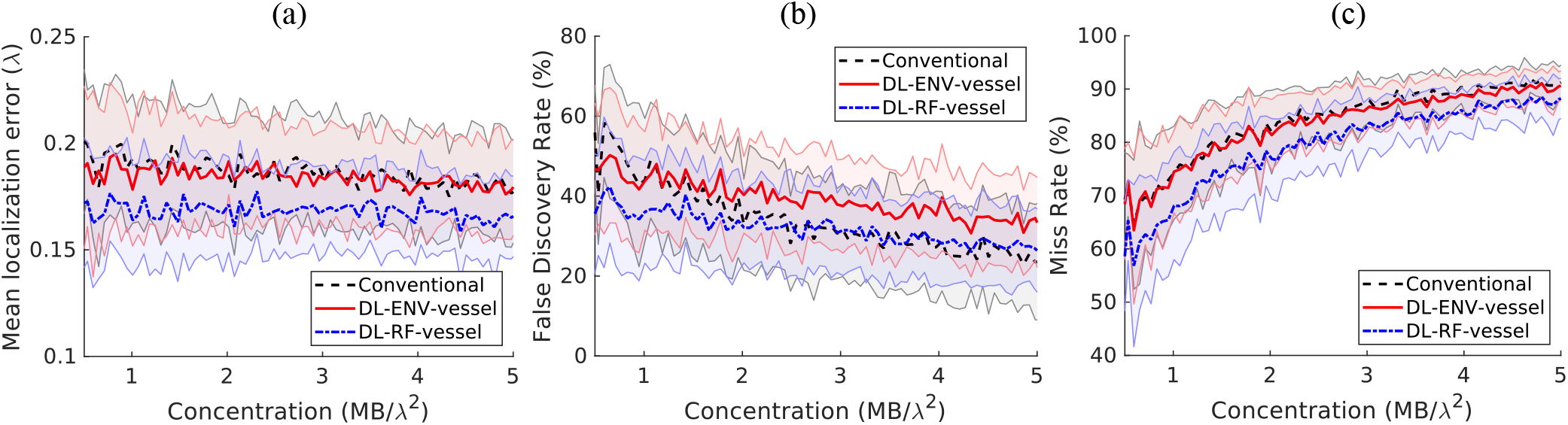
Localization performance on the simulation testing set, comparison between conventional localization, localization using DL-ENV and localization using DL-RF.

**Fig. 7.**
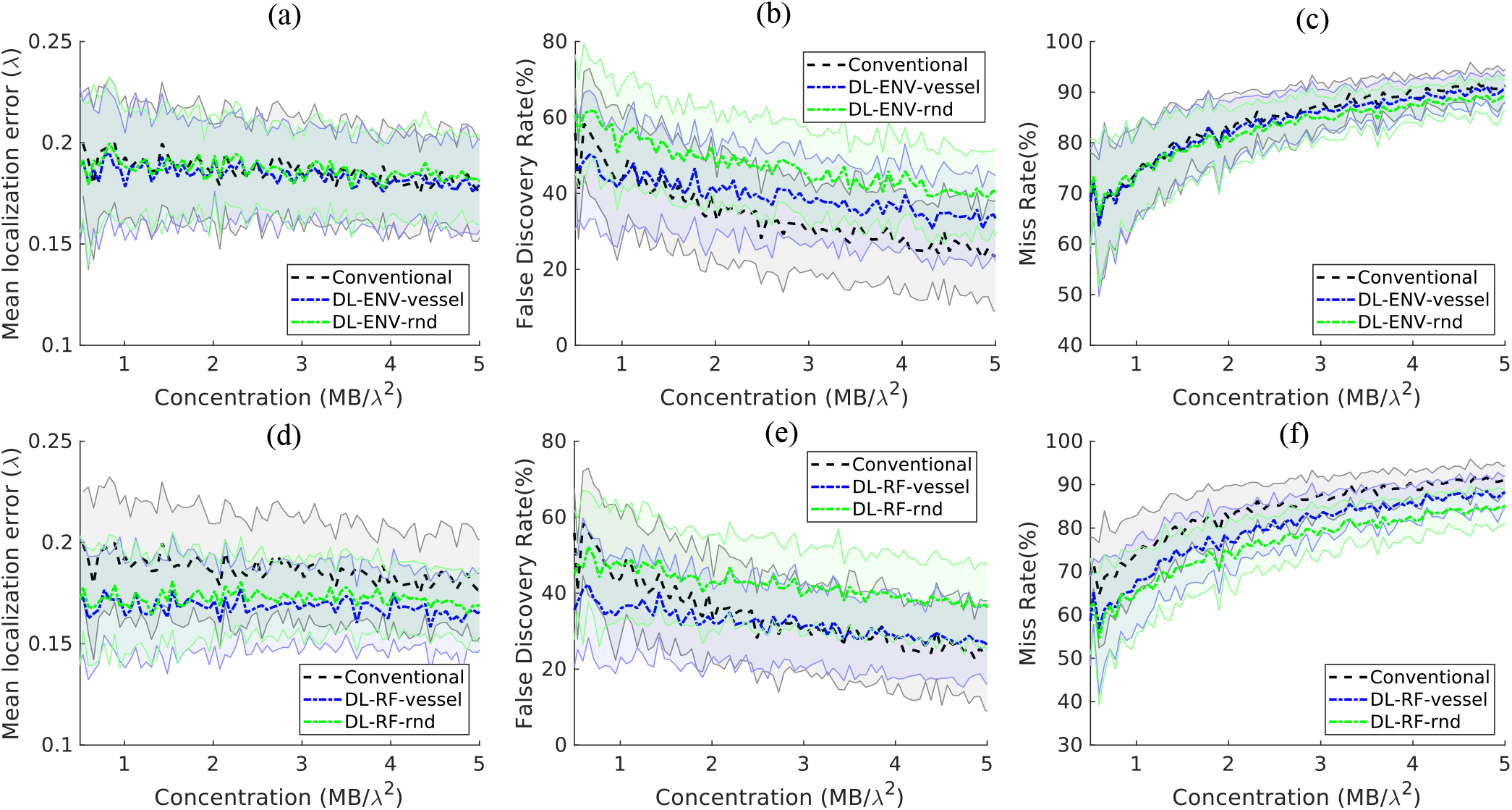
Localization performance on the simulation testing set, comparison between simulation training set with random MB distribution and MB within vessels. Note that the results presented in Fig. 6 were also included in Fig.7. The plots in Fig. 6 were rearranged to show comparison between conventional localization, localization using DL-ENV and localization using DL-RF.

Of note, beyond the concentration of 3MBs/λ^2^ or 500 MBs/mm^2^ (example shown in Fig. 8), the localization performance of all methods plateaued. The gain in MB localization performance using DL becomes very small compared to conventional localization. This is not surprising because at such high concentration, the task of recovering MB locations from ultrasound signal turned into an ill-posed problem. As it is not feasible to correctly localize individual MBs using a single frame of MB data with spatial information alone, regardless of the localization techniques being used, the previously introduced performance metric becomes less meaningful.

**Fig. 8.**
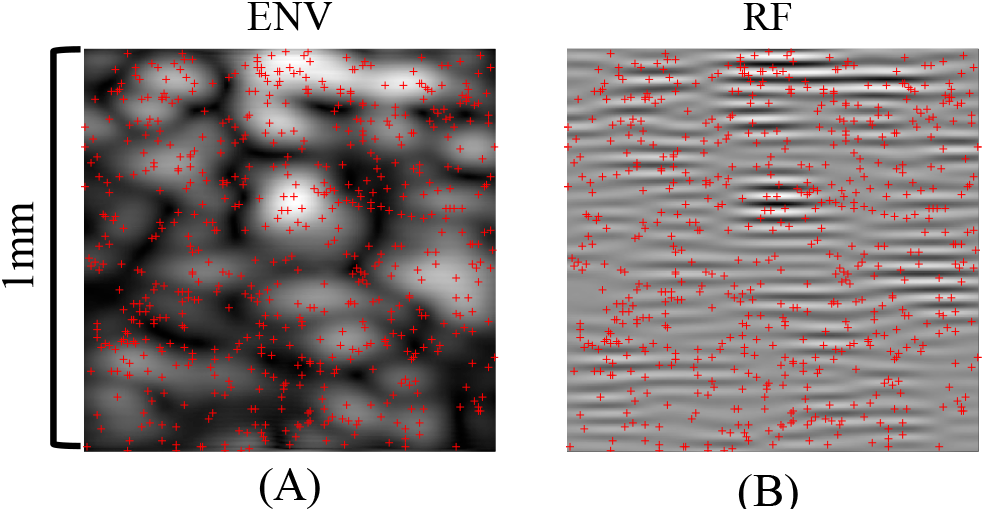
Ultrasound MB simulation where 625 MBs were present in the 1mm×1mm FOV. The ‘x’s mark the true MB locations. (A). ENV data. (B). RF data.

### C. Resolving synthetic simple vascular structure

When evaluated as a part of the ULM processing chain, the individual MB localization quality translates to the ability to reconstruct vessel structures with high fidelity. Therefore, we continued to investigate the quality of vessel maps reconstructed using each localization method. The evaluation was first performed on a set of synthetic data consisting of two vertical or horizontal pairs of vessels with a diameter of 5 pixels. Each pixel is 0.4928μm (approximately 0.064λ, the wavelength of an ultrasound imaging system with 20MHz center frequency). The separation between vessels was 2 pixels in the horizontal case, and 6 pixels in the vertical case. MBs were randomly placed within vessels to generate 200 frames of ultrasound data. ENV-trained network (DL-ENV), RF-trained network (DL-RF), and conventional localization were applied to each frame. Accumulating location of localized MBs over 200 frames results in the vessel maps shown in Fig. 9 (the horizontal case) and Fig. 10 (the vertical case). The profile of the reconstructed vessels averaged along the direction of the vessels were also shown in the two figures. For the horizontal vessels with a separation of 2 pixels (0.128 λ), DL-RF was able to produce a clean separation of the vessels. DL-ENV was also able to separate the vessels, but the separation was not as clear and only observed from the profile. In the vessel map generated using conventional localization, the two vessels were merged, shown by the absence of two separate peaks in Fig. 9C. When the separation was increased to 10 pixels (0.64 λ), all three methods where able to resolve the two vessels. However, DL-RF was able to produce the clearest boundary and cleanest nonvascular space in between vessels (Fig. 9B and D). Both DL-based localization methods were able to separate the two horizontal vessels with separation of 6 pixels (0.384 λ), where conventional localization failed to resolve two vessels. When the separation was increased to 10 pixels, all methods were able to resolve the vessels, with both DL-based methods producing cleaner boundaries than the conventional method and DL-RF slightly outperforming DL-ENV, which is consistent with the horizontal case. To summarize, both DL-based localization improved the ability to resolve closely positioned structures of ULM. Moreover, DL-RF was able to further improve the axial resolution due to its access to the information contained in the axial oscillation of the MB signal.

**Fig. 9.**
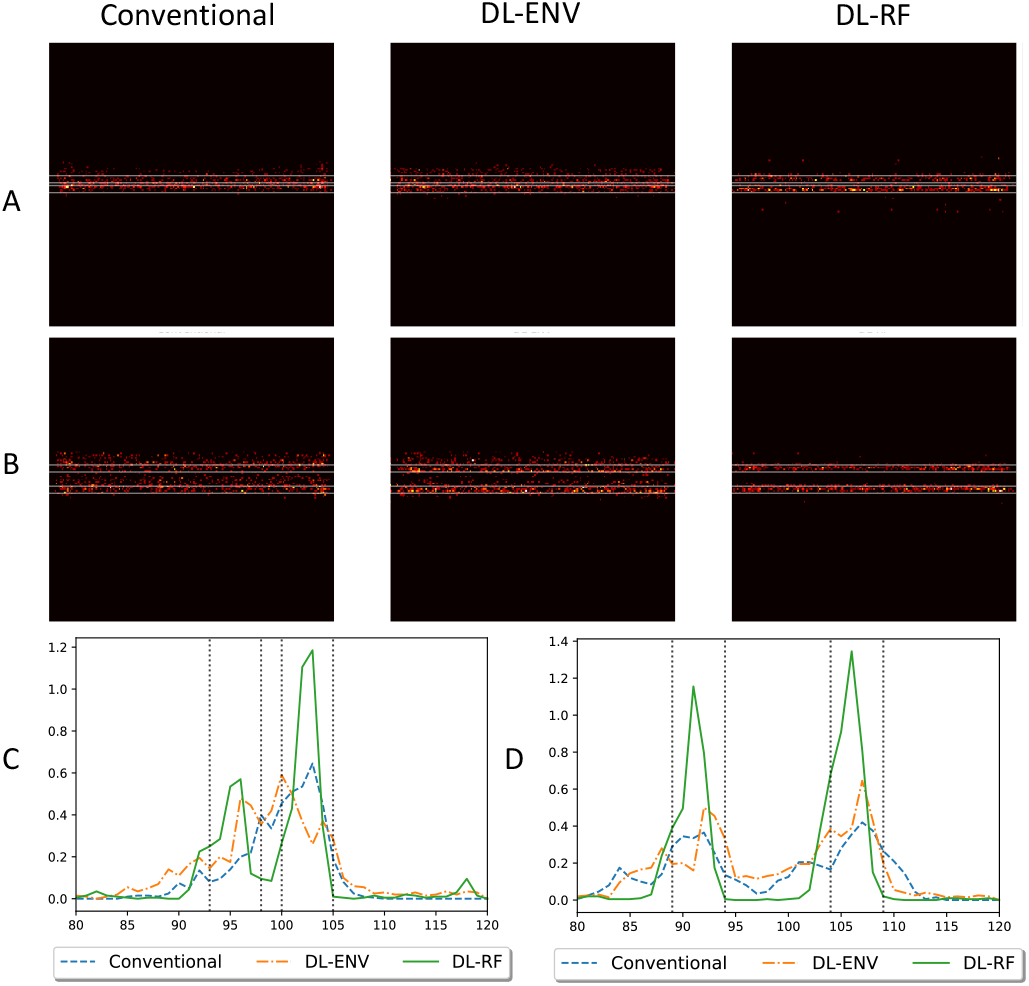
ULM reconstruction of two closely spaced simulation vessels. The vessels in A are axially separated by 2 pixels (0.9856 μm, 0.128 λ). The vessels in B are axially separated by 10 pixels (4.928 μm, 0.64 λ). C and D contain the profile of the reconstructed vessels in A and B, respectively.

**Fig. 10.**
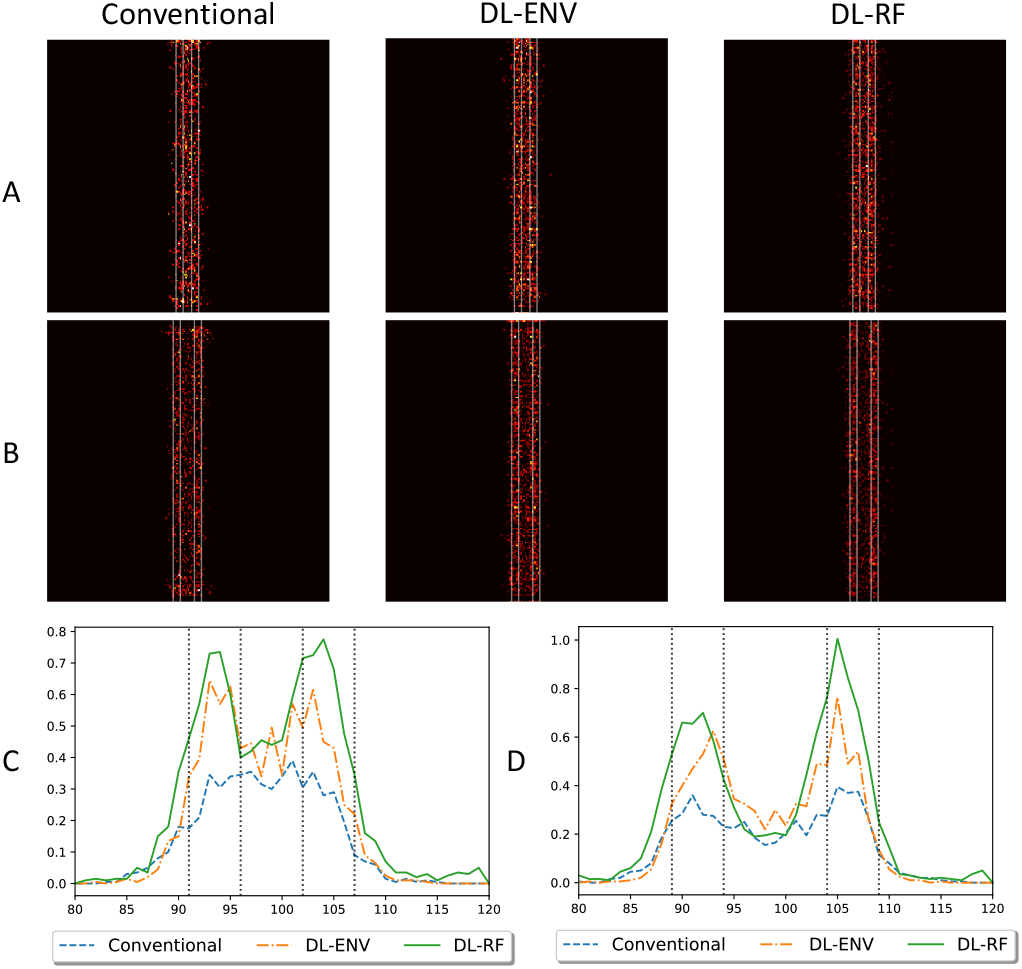
ULM reconstruction of two closely spaced simulation vessels. The vessels in A are laterally separated by 6 pixels (2.957 μm, 0.384 λ). The vessels in B are laterally separated by 10 pixels (4.928 μm, 0.64 λ). C and D contain the profile of the reconstructed vessels in A and B, respectively.

### D. MB localization performance using CAM vessel structures as testing data

In this part of the study, we tested the MB localization performance on simulation based on realistic CAM vessel structures. Only DL networks trained with group3 high concentration data (with CAM vessel structures) were used in this section of study. For test data generation, four 1.5mm×1.5mm regions of interest (ROIs) were sampled from the CAM optical images and converted to binary vessel maps using adaptive thresholding. For each vessel map, 1600 frames of ultrasound MB images were simulated using the same method described in Section II.A. The average MB concentration of the testing data was around 400 MB/mm^2^. Conventional and DL-based localizations were performed for each frame and accumulated across 1600 frames. Again, no MB tracking was conducted to better evaluate the localization performance for each method. Furthermore, the performance of localization methods was no longer evaluated based on the ability to precisely localize individual MBs. Two vessel-specific metrics were used: 1) the vessel false discovery rate (FDR) and 2) the vessel miss rate. Definitions for the vessel FDR and miss rate follow Eqs. (7) and (8), where false negatives (FN_vessel_) were defined as locations within vessels that were not covered by localization, false positives (FP_vessel_) were defined as locations outside vessels that were detected as localization, true positives (TP_vessel_) were defined as locations within vessels and covered by localization, and true negative (TN_vessel_) were as locations outside the vessels that do not have localizations.

Table II summarizes the localization results using 1600 frames of MB data. As shown in Table II, DL-RF localization was able to achieve both the lowest vessel miss rate and the lowest vessel FDR for all ROIs except for ROI 1 (although only 0.02% higher than the lowest FDR). DL-ENV had better miss rate performance than conventional localization (i.e., localize more MBs) but at the cost of elevated FDR.

**Table II.** MB Localization Performance Based on CAM Vessel Structures

|  |  | Conventional | DL-ENV | DL-RF |
| --- | --- | --- | --- | --- |
| ROI 1 | Miss Rate (%) | 26.52 | 22.67 | **18.11** |
|  | FDR (%) | **36.93** | 44.42 | 36.95 |
| ROI 2 | Miss Rate (%) | 25.83 | 23.33 | **18.54** |
|  | FDR (%) | 38.77 | 44.02 | **35.61** |
| ROI 3 | Miss Rate (%) | 26.36 | 21.05 | **18.31** |
|  | FDR (%) | 39.90 | 45.54 | **39.28** |
| ROI 4 | Miss Rate (%) | 26.89 | 22.86 | **19.65** |
|  | FDR (%) | 37.11 | 43.81 | **36.89** |

Fig. 11 shows the final localization results against the ground truth vessel map, where four different colors of pixels were used to represent FN_vessel_ (white), FP_vessel_ (blue), TP_vessel_ (red) and TN_vessel_ (black). Visually, it is apparent that the DL-RF method fills the vessel with the most TP_vessel_ red pixels while retaining a reasonable amount of erroneous blue pixels, indicating relatively fewer FP_vessel_ localizations than conventional DL-ENV. It is also worth noting that for both DL-based localization methods, the FP_vessel_ are most likely to occur just outside the edge of a vessel, while in the conventional localization, there are far more FP_vessel_ pixels in non-vessel regions. This result indicates that DL-based localization methods are less susceptible to randomly distributed noise commonly mis-localized as MBs than conventional localization techniques.

**Fig. 11.**
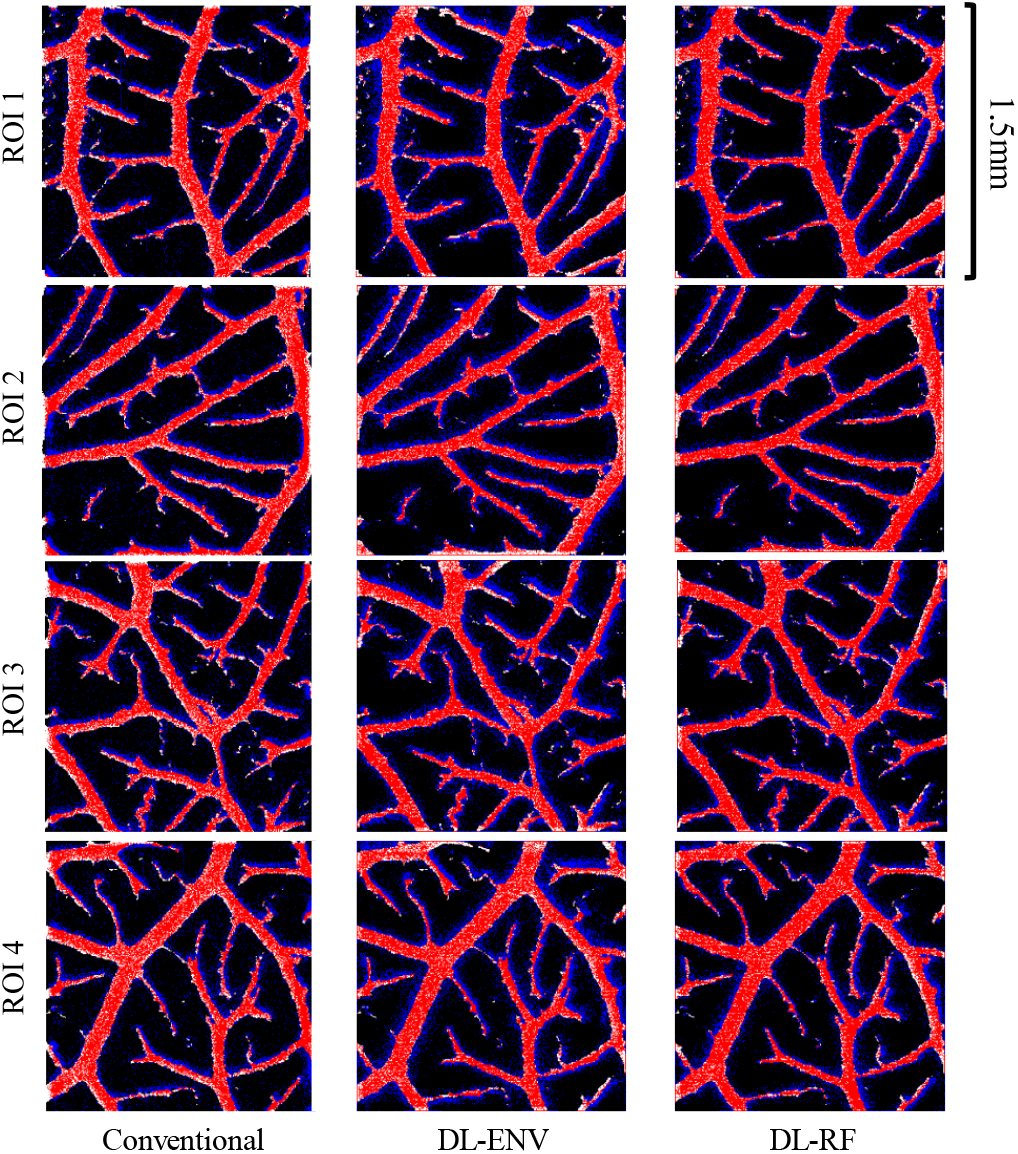
Difference maps of accumulated localization and ground truth vessel map for the simulation vessel dataset. Maps of the same ROI were arranged in the same row, whereas maps created using the same localization method were arranged in the same column. Red pixels represent TP_vessel_ localizations, blue pixels represent FP_vessel_ localizations, white pixels represent FN_vessel_ localizations, and black pixels represent TN_vessel_ localizations.

### E. In vivo study in CAM surface vessel

The proposed DL-based methods were then tested *in vivo* on a CAM surface vessel data. The generated microvessel density maps were validated against an optical image of the same area. Selected ROIs (Fig. 12) that provide best alignment between ultrasound and microscopy imaging were used for the comparison study. For each ROI, ground truth vessel masks were generated based on the optical image. The performance of each localization method was evaluated in terms of vessel saturation percentage (i.e., localized vessel pixels over total number of vessel pixels). Table III summarizes the 90% vessel saturation time for all the localization methods, including the MB separation method proposed in [25]. DL-RF localization was over 2X faster than the convention method to reach 90% vessel saturation, and over 20% faster than MB separation. This result is significant since MB separation is much more computationally expensive than the DL-based methods (excluding training).

**Fig. 12.**
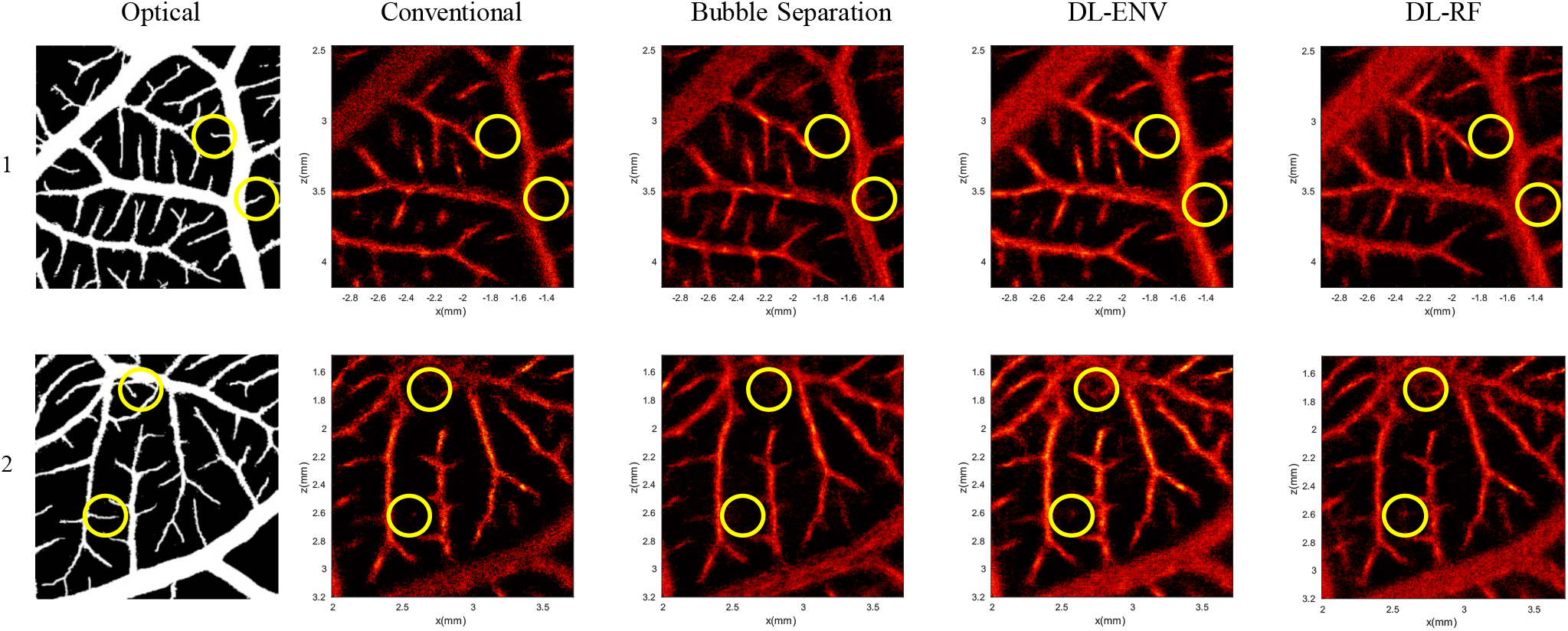
Comparison of different MB localization methods with optical ground truth for experimental CAM data. The circles denote example thin vessel segments where DL-based localization was better at localizing MB signals than the conventional techniques. The top row and bottom row are of two different ROIs. The first column is the binarized optical ground truth. The rest of the columns are the microvessel density map obtained using different localization methods.

**Table III.** Estimated Average Saturation Time

|  | Conventional | Bubble Separation | DL-ENV | DL-RF |
| --- | --- | --- | --- | --- |
| $t_{90\%}$ (s) | 4.458 | 2.397 | 2.526 | **1.958** |

Fig. 12 compares all localization methods against the optical ground truth. Both DL-based localization methods filled the large vessels better (e.g., higher MB count) than conventional localization and bubble separation, indicating a better MB isolation and localization performance at high MB concentrations. The circles in the figure highlight some thin vessel regions according to optical imaging, where DL-based methods had better performance than conventional methods. These results are consistent with the simulation results shown in Fig. 6 and Table II.

### F. Real-time high-resolution blood flow visualization

Fig. 13 (movie provided in Supplemental Videos 1 and 2) shows an example of using the proposed DL-based processing method to generate high-resolution, real-time visualization of blood flow *in vivo*. Thanks to the significantly enhanced spatial resolution of MB signal (Fig. 13B), one can directly accumulate multiple frames (e.g., 30 frames as shown in Fig. 13B and C) of the enhanced MB signal to produce high-resolution blood flow maps at a very fast rate with a very low computational cost (i.e., forward processing in the neural network). The accumulation alleviates the issue of sparse and weak MB signals within a single imaging frame and enhances the visualization of microvasculature. Longer accumulation time leads to better vessel delineation, at the cost of reduced temporal resolution. Fig. 14 (Supplemental Videos 3 and 4) demonstrates two small regions in sequential frames of the DL-enhanced display that captured some typical MB activities seen *in vivo*, including clusters of overlapping MB signal splitting and merging, which are difficult to visualize with conventional ULM.

**Fig. 13.**
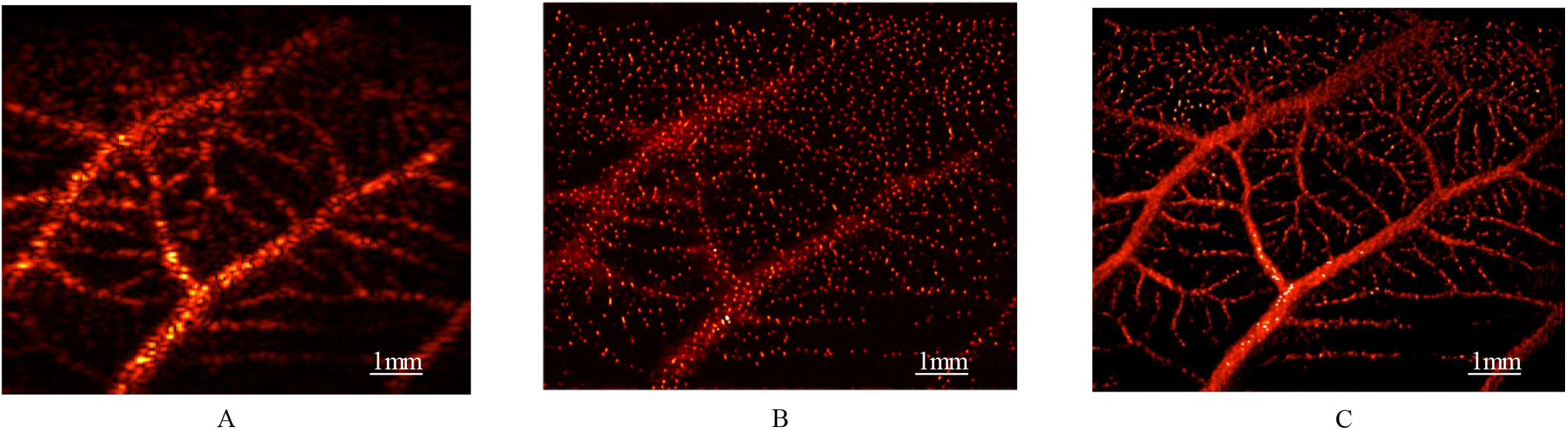
Example workflow for real-time high-resolution blood flow visualization using experimental CAM data. (A): the original B-mode input image (a single frame) of MBs in the CAM (movie provided in Supplemental Video 1). (B): DL-processed MB image using (A) as input. (C): A 30-frame accumulation of the DL-processed MB signals that demonstrates high-resolution blood flow visualization (movie provided in Supplemental Video 2).

**Fig. 14.**
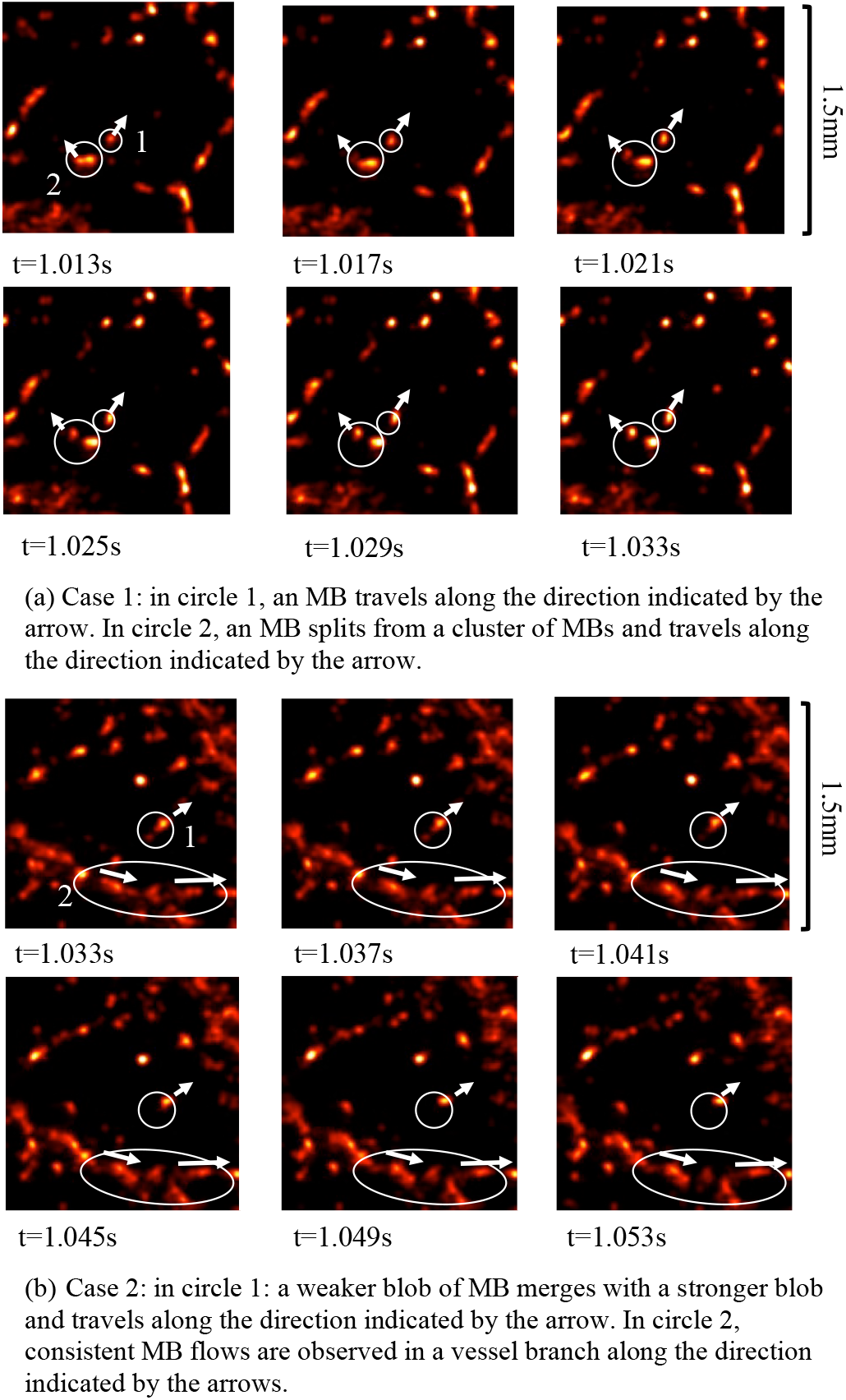
Two cases demonstrating the utility of the real-time high-definition MB imaging by DL processing on experimental CAM data.

### G. Computational cost

#### 1) ULM processing

Table IV summarizes the computational performance of different methods involved in this study, measured by the time consumption of the localization processing time on a single frame of CAM acquisition with size of 280 × 180 pixels (6.88 mm×8.83 mm). All algorithms were GPU-based, except for conventional localization that was implemented in both GPU and CPU for reference. The experiments were executed on a workstation running Ubuntu 18.04 operating system, with Intel^®^ Core™ i9-9820X @ 3.30GHz CPU (10 cores), NVIDIA^®^ GeForce^®^ RTX 2080 Ti GPU and 64GB of RAM. The conventional methods were implemented in MATLAB R2019b. The DL-based methods were implemented in Python 3.6 using PyTorch. DL-based methods in general have better computational performance than conventional. DL-ENV was the fastest, but the MB localization performance associated with DL-ENV was not as good as with DL-RF. DL-RF was able to achieve around 40% acceleration compared to localization with bubble separation while delivering better MB localization performance. The difference in computation time between DL-ENV and DL-RF was due to the IQ to RF conversion process, where the axial dimension was interpolated to 4 times the pixel size of the original IQ data. The input to the DL-ENV has the same spatial dimension with the original IQ data, which is 4 times smaller than the input to the DL-RF method.

**Table IV.** Computational Cost for Processing a Single Frame of CAM Data

| Method | Conventional<br>(CPU) | Conventional<br>(GPU) | Bubble<br>Separation | DL-ENV | DL-RF |
| --- | --- | --- | --- | --- | --- |
| Time(ms) | 601 | 130 | 304 | 77 | 168 |

#### 2) Real-time imaging feasibility of DL-enhanced, high-resolution blood flow imaging

The total time consumption for processing one frame of the MB data using a trained DL network and generating a new high-resolution vessel image is approximately 2.38 μs per pixel. The time consumption of Verasonics beamforming is ~40 ns per pixel for each compounding angle. Therefore, for a sequence that uses 9 compounding angles and acquires 100 × 100 pixel ultrasound data per frame, if ignoring the data transfer overhead, then the total processing time of each frame is (2.38 μs + 9 × 40 ns) × 10,000 pixels = 0.0274 s, which corresponds to a display frame rate of ~36 Hz.

For the processing time to satisfy a 10-Hz frame rate for realtime imaging, the processing time of each frame cannot exceed 0.1 s. Therefore, the input MB data may have up to 0.1 s ÷ (2.38 μs + 9 × 40 ns) = 36,496 ≈ 191 × 191 pixels. For a center frequency of 5 MHz (~0.3 mm pixel size), this image size corresponds to a 58.83 mm × 58.83 mm FOV. For a center frequency of 20 MHz (~0.077 mm pixel size), the image size corresponds to a 14.71 mm × 14.71 mm FOV. Note that these calculations did not factor in costs associated with clutter filters (e.g., SVD), although nonlinear imaging (e.g., pulse inversion, amplitude modulation) could be used in lieu of clutter filtering to extract MB signals to be fed into the neural network.

## IV. Discussion

In this study, we studied the performance of DL-based method for MB localization for ULM, using both envelope detected (ENV) and RF data under challenging high MB concentration scenario. We used Field-II to generate simulated training data sets both with random MB distributions and with realistic vascular structures obtained from *in vivo* CAM surface vessels. We found that having the vascular structure in the training data was beneficial for reducing the number of false positives of MB localization. Moreover, when underlying vascular structure information is available, imposing an additional structural constraint to further penalize localization in nonvascular regions can improve the quality of DL-based localization by producing cleaner nonvascular regions in the reconstruction. However, if realistic vascular structure is not available to use for simulation, completely randomly distribute MB would also provide serviceable result as the difference in mean localization error and miss rate were both minor. In general, DL-based localization methods showed better performance than conventional localization including MB separation. The performance of localization using RF data in both simulation and *in vivo* showed better performance over ENV since RF data contains both amplitude and phase information of the MB, consistent with the superior performance of RF than envelope reported for DL classification/regression [48]. We also identified three MB concentration ranges where the localization methods showed different types of behavior: for low concentration (under 0.2 MBs/λ^2^), more than half of the individual MBs can be localized with high confidence. Under moderate concentration (i.e. up to 1MBs/λ^2^), localization of individual MBs becomes unfeasible as concentration increases, but the detection of the centroids of multiple clustered MBs can still be used for ULM reconstruction, and the increased concentration provides faster saturation. Finally, the performance of both conventional and DL-based localization plateaus as MB concentration reaches extreme highs (e.g., > 3 MB/λ^2^). Increasing MB concentration is no longer profitable, even with a more robust localization mechanism. Under high MB concentrations, it is possible that information regarding individual MB locations no longer exists in the RF data due to significant MB signal overlap (i.e., ultrasound wave interference cannot be recovered using only spatial information). In order to overcome this limitation, modifying the DL model to also incorporate temporal information of MBs becomes necessary to further improve DL-based MB localization performance.

For *in vivo* ULM imaging, adjusting the concentration of contrast agents in the blood stream can be challenging, especially for imaging under clinical settings. Even if a lower MB concentration can be used to facilitate more robust MB localization, MB concentration can still be very high within large vessels and arteries. Moreover, *in vivo* animal and human imaging are in general more challenging than CAM due to deeper imaging depth, tissue motion, and other sources of noise such as multi-path reverberation and phase aberration. Nevertheless, based on the promising results shown in this study, future studies targeting the development of a more robust DL-based MB localization method that can account for these challenges are warranted.

There are some limitations in this study: First, although optical images of the CAM surface vessel were used as ground truth for evaluating the performance of MB localization in vivo, it was difficult to achieve a perfect registration between ultrasound and optical imaging FOVs. The quality of the ground truth obtained from optical images was also affected by the quality of image segmentation as well as the resolution limit of the optical imaging system. As a result, the evaluation metric may not deliver an entirely accurate evaluation of the MB localization performance *in vivo*; second, simulations such as Field II cannot perfectly account for the variability of PSF in the real image system. Also, Field II may not appropriately simulate nonlinear MB response subject to ultrasound. Therefore, the training of the neural networks was not optimal and can be further improved. One solution to this issue is to carefully align ultrasound FOV with optical imaging and simultaneously acquire ultrasound and optical MB data (with MB fluorescently labeled), which can then be used for training. However, the amount of training samples that can be obtained using this method will be far less than from using simulation. Therefore, we posit that simulation will still serve an important role to initialize DL training for MB localization, and experimental ground-truth data can be used for fine-tuning the network to boost its performance.

ULM is computationally expensive, and therefore any improvement to any segment of the processing chain contributes to the reduction of overall processing time. In this paper, we demonstrated the computational advantage of DL-based MB localization over conventional localization techniques. Our analysis of computational time did not include the clutter filtering and motion correction steps because they were performed for both DL-based and conventional localization techniques. The computational cost of these additional processing steps can significantly affect the real-time capability of the DL-based localization technique. The additional processing time can be optimized by utilizing more efficient algorithms, such as DL-based clutter suppression [49] [50]. Moreover, we did not conduct further downstream processing of the MB locations such as MB tracking in this study, which can be more computationally expensive than localization. Nevertheless, DL should also be capable of tracking MB locations using recurrent neural network architectures such as the long short-term memory [51]. Finally in this paper, we proposed a real-time feasible high-resolution vessel visualization method that directly uses DL-enhanced MB signals to display vessels. Although the high-resolution display allows one to visually perceive the blood flow dynamics, the image does not contain quantitative flow speed information that can be obtained with conventional ULM. However, direct inference of MB velocity may be possible by training the DL network with spatial-temporal sequences of MB movement, which ultimately allows real-time display of blood flow speed at high spatial resolution.

## V. Conclusion

In this paper, we studied the performance of DL-based MB localization method for ULM for increasing MB concentration. A U-Net style convolutional neural network was trained with Field-II simulated ultrasound data (both envelope detected and RF data) based on *in vivo* CAM surface vessel structure. The neural network was able to sharpen ultrasound MB signal, so that overlapping and distorted MB signal were able to be better localized. Under low concentration, DL-based methods were able to more precisely localize individual MBs. As concentration increased, the behavior of all localization methods changed to localizing speckles rather than individual MB. The DL-based method was still able to improve the localization accuracy and reduce saturation time by localizing more MB clusters in each frame of ultrasound acquisition. Furthermore, the DL-processed ultrasound data can be utilized to achieve real-time high-resolution visualization of blood flow dynamics. The proposed method can be further improved by developing a training data set with better ground truth and incorporating temporal information of microbubbles in training.

